# Bayesian Inference of Species Networks from Multilocus Sequence Data

**DOI:** 10.1101/124982

**Authors:** Chi Zhang, Huw A. Ogilvie, Alexei J. Drummond, Tanja Stadler

## Abstract

Reticulate species evolution, such as hybridization or introgression, is relatively common in nature. In the presence of reticulation, species relationships can be captured by a rooted phylogenetic network, and orthologous gene evolution can be modeled as bifurcating gene trees embedded in the species network. We present a Bayesian approach to jointly infer species networks and gene trees from multilocus sequence data. A novel birth-hybridization process is used as the prior for the species network, and we assume a multispecies network coalescent (MSNC) prior for the embedded gene trees. We verify the ability of our method to correctly sample from the posterior distribution, and thus to infer a species network, through simulations. To quantify the power of our method, we reanalyze two large datasets of genes from spruces and yeasts. For the three closely related spruces, we verify the previously suggested homoploid hybridization event in this clade; for the yeast data, we find extensive hybridization events. Our method is available within the BEAST 2 add-on `SpeciesNetwork`, and thus provides an extensible framework for Bayesian inference of reticulate evolution.

## 1 Introduction

Hybridization during speciation is relatively common in animals and plants (Mallet, 2005, 2007). However, when reconstructing the evolutionary history of species, typically non-reticulating species trees are inferred (Guindon et al., 2010; Stamatakis, 2014; Drummond and Bouckaert, 2015; Ronquist et al., 2012), and the potential for hybridization events is ignored.

To account for the distribution of evolutionary histories of genes inherited from multiple ancestral species, the multispecies coalescent model (Rannala and Yang, 2003; Liu et al., 2009) was extended to allow reticulations among species, named multispecies network coalescent (MSNC) model (Yu et al., 2014). Orthologous genes are modeled as gene trees embedded in the species network. The MSNC model accounts for gene tree discordance due to incomplete lineage sorting and reticulate species evolution events, such as hybridization or introgression. There have been computational methods developed based on the MSNC to infer species networks using maximum likelihood (Yu et al., 2014; Yu and Nakhleh, 2015; Solís-Lemus and Ané, 2016) and Bayesian inference (Wen et al., 2016). These methods use gene trees inferred from other resources as input. Due to the model complexity, applying the MSNC model in a full Bayesian framework, i.e., to infer the posterior distribution of species network and gene trees directly from the multilocus sequence data, is challenging. Recently Wen and Nakhleh (2017) have developed a Bayesian method that can co-estimate species networks and gene trees from multilocus sequence data, but a process-based prior for the species network is still lacking. Their method also integrates over all possible gene tree embeddings at each MCMC step, which means that the estimated histories of individual gene trees within the species network are not available for subsequent analysis, and the method does not co-estimate base frequencies or substitution (transition and transversion) rates.

In this paper, we present a Bayesian method to infer ultrametric species networks jointly with gene trees and their embeddings from multilocus sequence data. Our method assumes a birth-hybridization model for the species network, the MSNC model for the embedded gene trees with analytical integration of population sizes, and employs novel MCMC operators to sample the species network and gene trees along with associated parameters. It is able to use the full range of substitution models implemented in BEAST 2 (Bouckaert et al., 2014), including models with gamma rate variation across sites (Yang, 1994).

## 2 New Approaches

In this section, we specify our approach to sample from the posterior distribution of species networks and gene trees, given a multilocus sequence alignment. First we derive the unnormalized posterior distribution. Then we introduce operators to move through the space of species networks, the space of gene trees, and finally to update the gene tree embeddings within species networks.

### 2.1 The posterior distribution of species networks and gene trees

#### 2.1.1 The probability density of a species network

The birth-hybridization process provides a prior probability for a given species network Ψ (Fig. 1). The process starts from *t*_0_ (time of origin) in the past with a single species. A species gives birth to a new species with a constant rate λ (speciation rate), and two species merge into one with a constant rate *v* (hybridization rate). That is, at the moment of *k* species, the speciation rate is *kλ*, the hybridization rate is 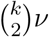, and the waiting time to the next event is an exponential distribution. The process ends at time 0 (the present).

**Figure 1:**
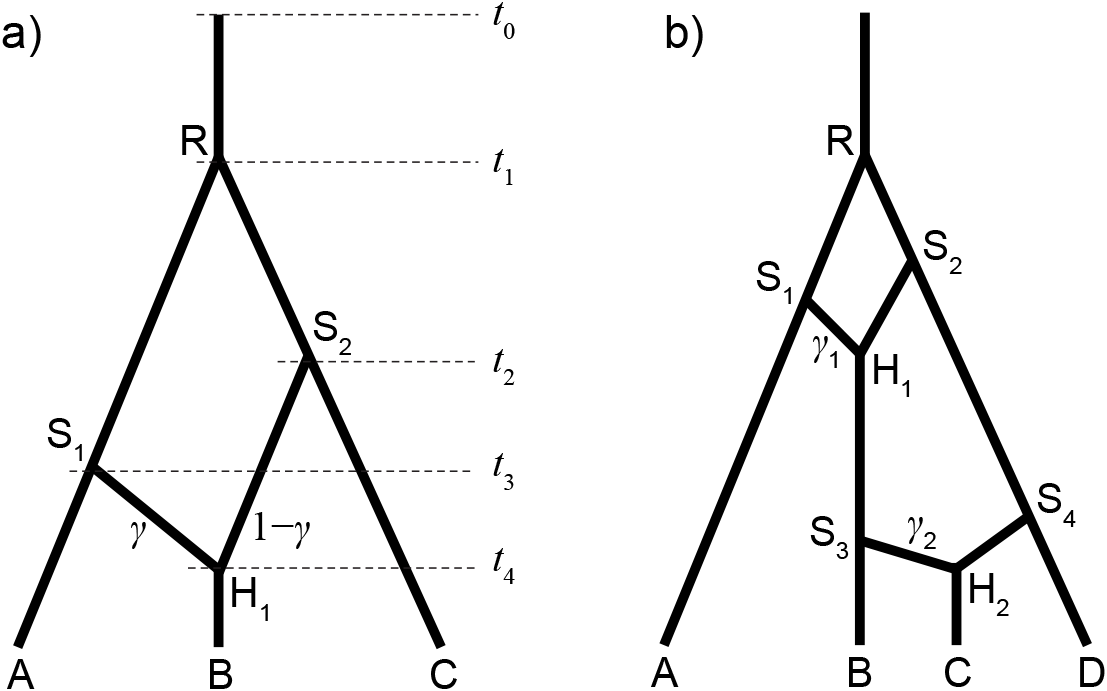
a) A species network with 3 tips, 3 bifurcations, and 1 reticulation. The inheritance probability at branch *S*_1_*H*_1_ is γ, and that at *S*_2_*H*_1_ is 1 − γ. b) Another network with 4 tips and 2 reticulations, with γ_1_ and γ_2_ associated with *S*_1_ *H*_1_ and *S*_3_ *H*_2_, respectively.

The probability density of a species network Ψ with *n* extant species descending from *n* − 1 + *m* speciation events and *m* hybridization events, and these events happening at time *t*_1_ > *t*_2_ >…> *t_n+2m-1_*, conditioned on *t*_0_, λ and *v*, is,

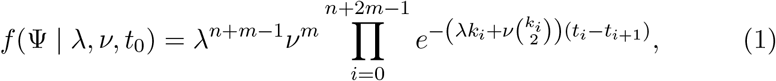

where *k_i_* is the number of lineages within time interval (*t*_*i*_, *t*_*i*+1_) and *t*_*n*+2*m*_ = 0 is the present time. For the network shown in Figure 1a, the probability density of the species network is

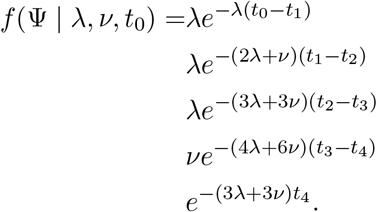

In our Bayesian analysis, the parameters λ, *v*, and *t*_0_ can be assigned hyperpriors.

#### 2.1.2 The probability of the sequence data given the gene trees

Assuming complete linkage within each locus, the probability of the data *D* = {*D*_1_, *D*_2_,…,*D*_l_} given gene trees *G* = {*G*_1_, *G*_2_,…,*G*_l_} is the product of phylogenetic likelihoods (Felsenstein, 1981) at individual loci:

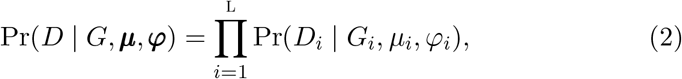

where *G_i_* is the gene tree with coalescent times, *μ_i_* is the substitution rate per site per time unit, and φ_*i*_ represents the parameters in the substitution model (e.g., the transition-transversion rate ratio κ in the HKY85 model (Hasegawa et al., 1985)), at locus *i* (*i* = 1,…,L).

There are two sources of evolutionary rate variation: across gene tree lineages at the same locus and across different gene loci. In the strict molecular clock model (Zuckerkandl and Pauling, 1965), μ is the global clock rate, i.e., no rate variation across gene lineages at each locus. To extend to a relaxed molecular clock model (e.g., Thorne and Kishino, 2002; Drummond et al., 2006; Lepage et al., 2007; Rannala and Yang, 2007), the molecular clock rate is variable across gene lineages following certain distributions with μ as the mean. To account for rate variation across genes, gene-rate multipliers {*m*_1_, *m*_2_,…,*m*_L_} are constrained to average to 1.0 (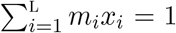, where *x_i_* is the proportion of sites in locus *i* to the total number of sites). Then the substitution rate at locus *i* is *μ_i_ = μm_i_*. Thus, when multiplying the gene tree lineages in *G_i_* by *μ_i_*, all the branch lengths are then measured by genetic distance (substitutions per site).

The gene-rate multipliers are assigned a flat Dirichlet prior. The average substitution rate (clock rate) μ can be either fixed to 1.0 such that branch lengths are measured by genetic distance, or assigned an informative prior to infer branch lengths measured in absolute time.

#### 2.1.3 The probability density of the gene trees given a species network

The gene trees *G* = {*G*_1_, *G*_2_,…,*G*_L_} are embedded in the species network under the multispecies network coalescent (MSNC) model (Yu et al., 2014) (Fig. 2). Hybridizations or horizontal gene transfers are modelled by reticulations in the species network. The effective population sizes *N* = {*N*_1_, *N*_2_,…, *N*_B_} are assumed to be identically and independently distributed (i.i.d.) for each of the b branches in Ψ, while each locus has the same effective population size *N_i_* at branch *i* (*i* = 1,…,B). For each locus *j*, the number of coalescences of gene tree *G_j_* within branch *b* of is denoted by *k_jb_*, and the number of lineages at the tipward end of *b* is denoted by *n_jb_*, thus the number of lineages at the rootward end of *b* is *n_jb_ – k_jb_*. The *k_jb + 1_* coalescent time intervals between the tipward and rootward of branch *b* are denoted by *c_jbi_* (0 ≤ *i* ≤ *k_jb_*). *p_j_* is the gene ploidy of locus *j* (e.g., 2 for autosomal nuclear genes and 0.5 for mitochondrial genes in diploid species). **γ** = {γ_1_, …,γ_H_} are the inheritance probabilities, one per reticulation node in Ψ. For each lineage of *G_j_* traversing the reticulation node *H_h_* backward in time, with probability *γ_h_* it goes to the parent branch associated with that inheritance probability, and to the alternate parent branch with probability 1 − *γ_h_*. The corresponding number of traversing lineages are denoted by *u_jh_* and *v_jh_* respectively.

**Figure 2:**
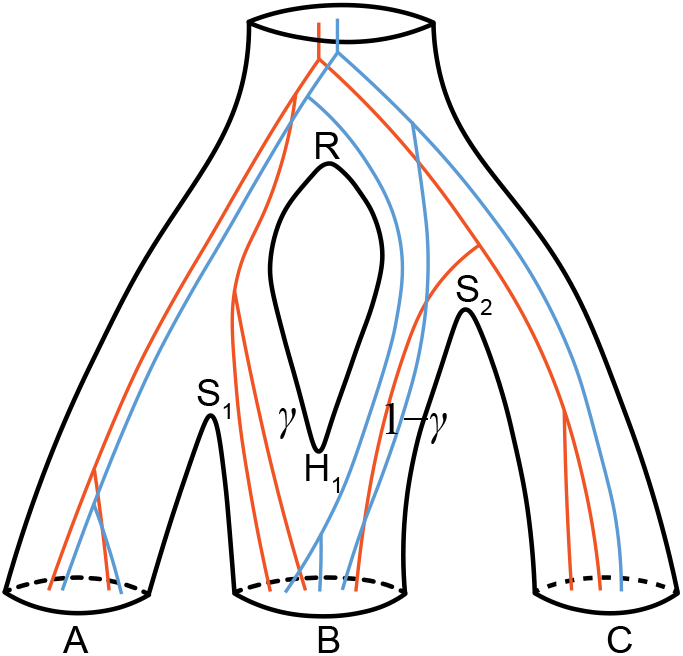
Two gene trees embedded in the species network of Fig. 1a. There are 2 samples from species *A*, 3 samples from *B*, and either 1 or 2 samples from *C*. For each gene tree lineage traversing the reticulation node *H*_1_ backward in time, it goes to the left population with probability *γ*, and to the right with probability 1 − *γ*.

The coalescent probability of the gene trees *G* in species network Ψ with time being measured in calendar units is thus:

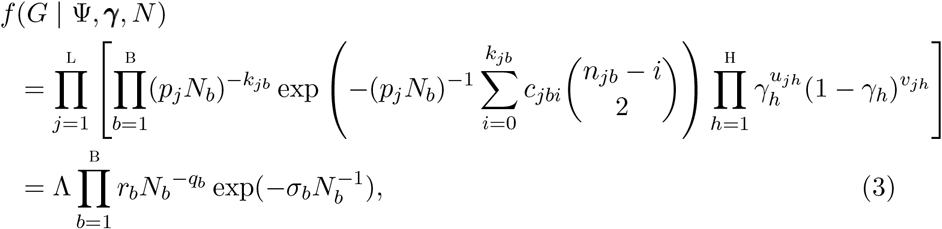

where 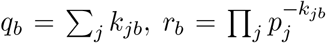, 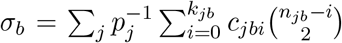 and 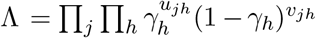. When there is no reticulation in the species network (i.e., it is a species tree), then Λ = 1 and Equation 3 is equivalent to Equation 2 in Jones (2017).

Note here, when time is measured by genetic distance, we use *θ_b_ = N_b_μ* as the population size parameter of branch *b*, and *τ_i_ = t_i_ μ* as the height of node *i*. The prior for **γ** can be any distribution on [0, 1], we use throughout *f(γ_h_) ~ U*(0, 1). In the next section, we discuss how to integrate out the population sizes, which will improve computational speed.

#### 2.1.4 Integrating out the population sizes analytically

Equation 3 has the form of unnormalized inverse gamma densities. The population sizes N can be integrated out through the use of i.i.d. inverse-gamma(*α, β*) conjugate prior distributions (Jones, 2017; Hey and Nielsen, 2007), that is,

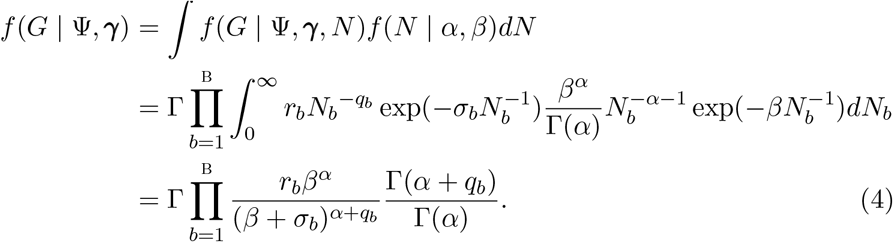

The symbolic notations follow Equation 3.

#### 2.1.5 The joint posterior distribution

The joint posterior distribution of the parameters is

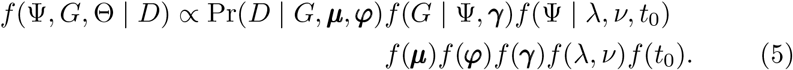

Here Θ represents (***μ, φ, γ**, λ, v, t*_0_).

### 2.2 MCMC operators for the species network

#### 2.2.1 Node slider

The node-slider operator only changes the node heights of the species network, not the topology. It selects an internal node or the origin randomly, then proposes a new height centered at the current height according to a normal distribution: *t'* | *t* ~ *N*(*t*,*σ*^2^), where σ is a tuning parameter controlling the step size. The lower bound is the oldest child-node height, the upper bound is the youngest parent-node height (except for the origin, Fig. 3). If the proposed value is outside this range, the excess is reflected back into the interval. Note that for the origin, if the proposed height is outside the range of its prior, this move is aborted. A variation of this operator can use a uniform proposal instead of the normal proposal: *t'* | *t* ~ *U* (*t* − *w*/2, *t* + *w*/2), where *w* is the window size. The proposal ratio is 1.0 in both cases.

**Figure 3:**
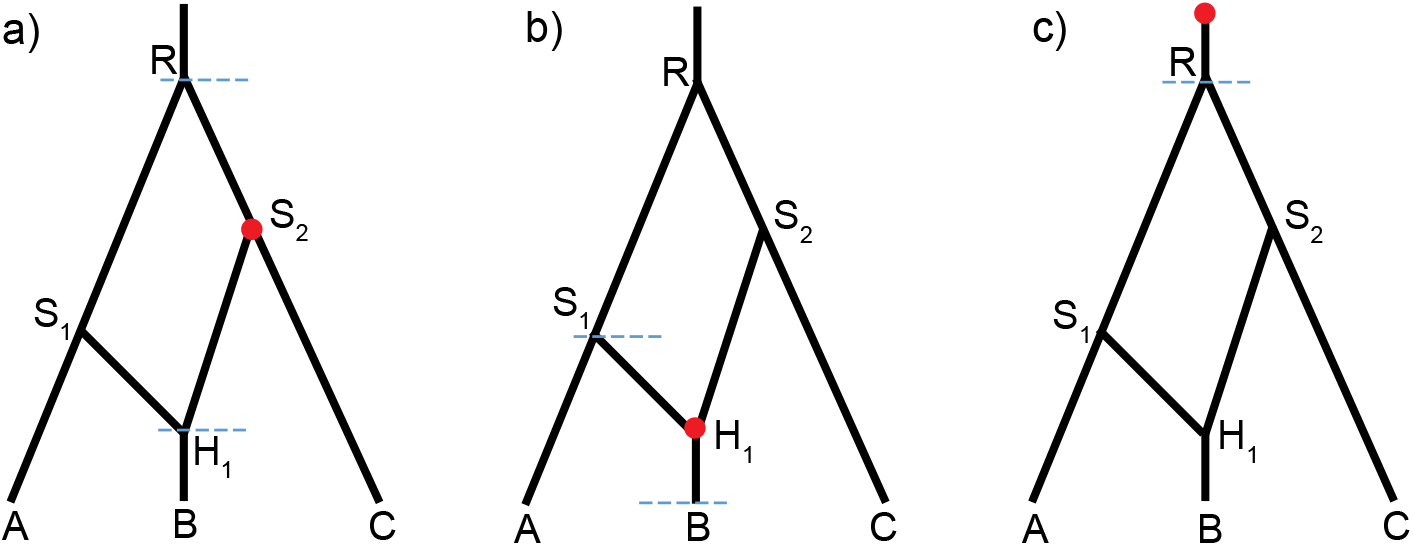
Three cases when the node-slider operator is applied: a) a bifurcation node *S*_2_ is selected; b) the reticulation node *H*_1_ is selected; c) the origin is selected. The dashed lines are the lower and upper bounds for changing its height (only the lower bound is applicable in c)). For the node-uniform operator, a) and b) apply but c) does not.

#### 2.2.2 Node uniform

The node-uniform operator also changes the internal-node heights of the species network while keeping the topology. It selects an internal node randomly, then proposes a new height uniformly between the lower and upper bounds (Fig. 3ab). The lower bound is the oldest child-node height, the upper bound is the youngest parent-node height. The proposal ratio is 1.0. Unlike node slider, this operator does not change the time of origin. A separate operator for the origin, such as multiplier or scaler, can be coupled to update all the node heights.

#### 2.2.3 Relocate Branch

The relocate-branch operator can change the topology, but keeps the number of reticulations in the species network constant. It first selects an internal node at random. If the selected node is a bifurcation node, the rootward end of either its child branches is selected (Fig. 4a); if the selected node is a reticulation node, the tipward end of either its parent branches is selected (Fig. 4b). Then the selected branch is detached at the side of the selected node, and a destination branch to be attached is chosen randomly from all possible candidate branches (including the original position). A new height of the selected node is proposed uniformly between the heights of the two ends of the destination branch (*v*' and *u*' in Fig. 4). When the relocated branch has a bifurcation node at one end and a reticulation node at the other end, the candidate branches include all the remaining branches, and the reticulation direction can be changed depending on the proposed new height (Fig. 4b). When the relocated branch has the same type of nodes at both ends and the resulted network is invalid, the move is aborted. For example, moving the rootward end lower than the tipward end if the two ends are both bifurcation nodes, or moving the tipward end higher than the rootward end if the two ends are both reticulation nodes, will result in an invalid network. We denote with v and u the lower and upper bounds of the backward move. The proposal ratio is (*u*' − *v*')/(*u* − *v*).

**Figure 4:**
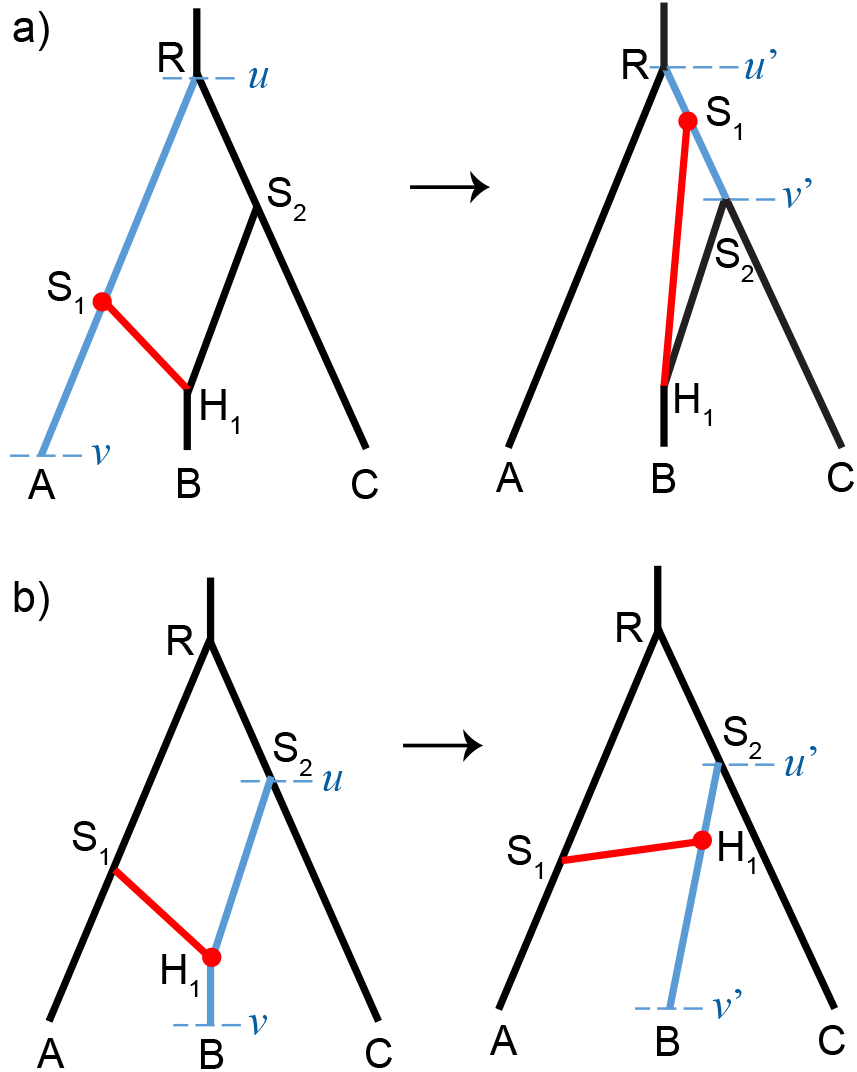
Two cases when the relocate-branch operator is applied. a) A bifurcation node *S*_1_ is selected, and branch *S*_1_ *H*_1_ is relocated to attach to *RS*_2_. b) A reticulation node *H*_1_ is selected, and branch *S*_1_ *H*_1_ is still attaching to *S*_2_*B* with flipped reticulation direction. The lower and upper bounds of proposing the new attaching point are *v*' and *u*', and the corresponding bounds of the backward move are *v* and *u*.

#### 2.2.4 Add- and delete-reticulation

The add-reticulation and delete-reticulation operators are reversible-jump MCMC (rjMCMC) proposals that can add and delete a reticulation event respectively.

In the add-reticulation operator, a new branch is added by connecting two randomly selected branches with length *l*_1_ and *l*_2_ (Fig. 5). The same branch can be selected twice so that *l*_1_ = *l*_2_ (Fig. 5b). Then three values *w*_1_, *w*_2_ and *w*_3_ are drawn from *U*(0, 1). One attaching point cuts the branch length *l*_1_ to *l*_11_ = *l*_1_ *w*_1_ (and thus *l*_12_ = *l*_1_ (1 − *w*_1_)); the other attaching point cuts the branch length *l*_2_ to *l*_21_ = *l*_2_ *w*_2_ (and thus *l*_22_ = *l*_2_ (1 − *w*_2_)). Analogously, if we select the same branch twice, the attachment times of the new branch are *l*_1_ *w*_1_ and *l*_1_ *w*_2_. An inheritance probability γ = *w*_3_ is associated to the new branch. We will operate on the inheritance probability γ of this added branch, while the inheritance probability of the second reticulation branch (i.e., 1 − γ) changes accordingly. We denote *k* as the number of branches in the current network, and *m* as the number of reticulation branches (parent branches of the reticulation nodes) in the proposed network. The Hastings ratio is then (1/*m*)/[(1/*k*)(1/*k*) × 1 × 1 × 1] = *k*^2^/*m*. The Jacobian is 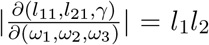. Thus the proposal ratio of add-reticulation is *l*_1_*l*_2_ *k*^2^/*m*.

**Figure 5:**
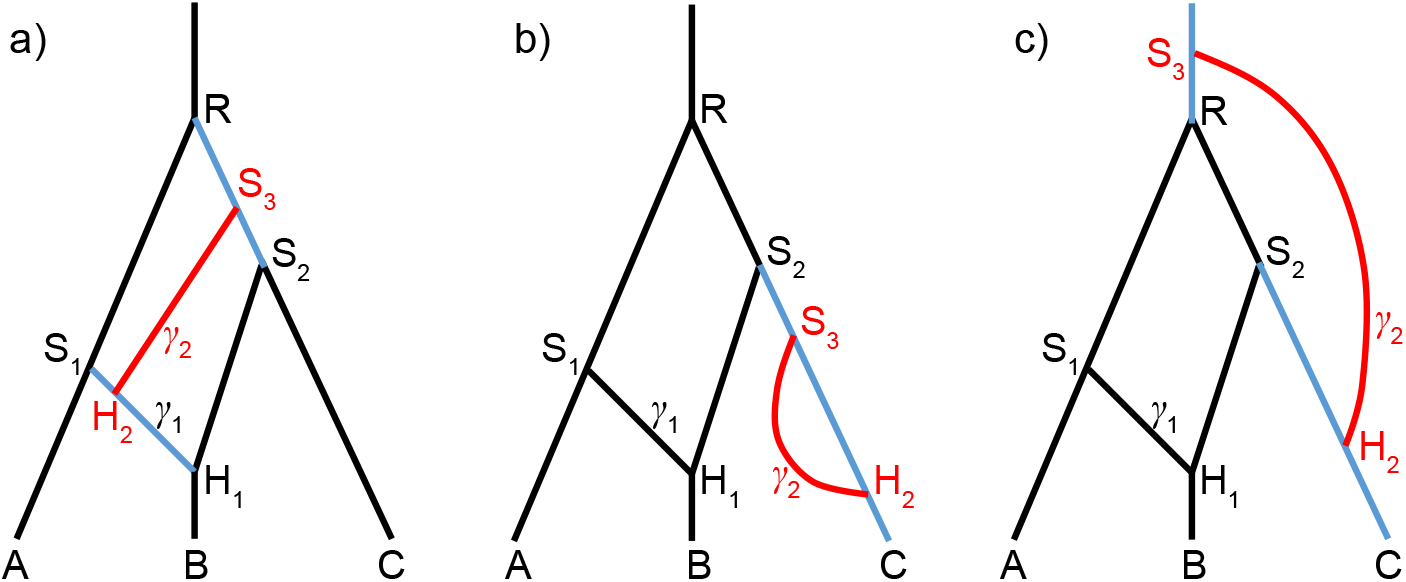
Three cases when the add-reticulation operator is applied. The number of branches in the current network (i.e., the network without the red branch) is *k* = 8. The probability of selecting the illustrated branches (in blue) is 1/*k*^2^. The number of reticulation branches in the proposed network is *m* = 4. In the reverse move, delete-reticulation, the probability of selecting the added branch (in red) is 1/*m*. a) Branches *S*_1_ *H*_1_ and *RS*_2_ are selected and a new branch *S*_3_ *H*_2_ is added together with γ_2_. The length of *S*_1_*H*_1_ is *l*_1_ = *l_S_1_H_1__*, and that of *RS*_2_ is *l*_2_ = *l_RS_2__*. In the delete-reticulation move, if *H*_1_ *H*_2_ is selected, the operator is aborted. b) The same branch *S*_2_*C* is selected twice. *l*_1_ = *l*_2_ = *l*_S_2_C_, *l*_11_ = *l*_S_2_S_3__, *l*_21_ = *l*_S_2_H_2__. c) The root branch and *S*_2_*C* are selected. *S*_3_ becomes the new root.

In the delete-reticulation operator, a random reticulation branch together with the inheritance probability γ is deleted (Fig. 5). Joining the singleton branches at each end of the deleted branch, resulting in two branches with length *l*_1_ and *l*_2_ completes the operator (*l*_1_ = *l*_2_ when forming a single branch, Fig. 5b). If there is no reticulation, or the selected branch is connecting two reticulation nodes, the move is aborted. For example in Figure 5a, deleting reticulation branch *H*_1_ *H*_2_ will result in an invalid network. We denote *k* as the number of branches in the proposed network, and *m* as the number of reticulation branches in the current network. The proposal ratio of delete-reticulation is *m*/(*k*^2^*l*_1_*l*_2_).

#### 2.2.5 Inheritance-probability uniform

The inheritance-probability uniform operator selects a reticulation node randomly, and proposes a new value of the inheritance probability *γ' ~ U*(0, 1). The proposal ratio is 1.0.

#### 2.2.6 Inheritance-probability random-walk

The inheritance-probability random-walk operator selects a reticulation node randomly, and applies a uniform sliding window to the logit of the inheritance probability γ, that is *y*' | *y ~ U (y-w/2,y+w/2)*, where *y* = logit(*γ*) = log(*γ*) − log(1 − *γ*). Since the proposal ratio for the transformed variable *y* is 1.0, and 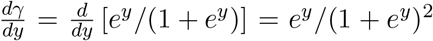, the proposal ratio for the original variable *γ* is 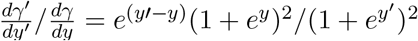.

### 2.3 MCMC operators for gene trees

The standard tree operators in BEAST 2 (Bouckaert et al., 2014) are applied to update the gene trees, including the scale, uniform, subtree-slide, narrow- and wide-exchange, and Wilson-Balding (Wilson and Balding, 1998). The scale and uniform operators only update the node heights without changing the tree topology, while the other operators can change the topology (Drummond and Bouckaert, 2015). The species network is kept unchanged when operating on the gene trees, and vice versa.

### 2.4 MCMC operator for the gene tree embedding

The gene trees must be compatibly embedded in the species network (Fig. 2). When a new gene tree is proposed using one of the tree operators, the rebuild-embedding operator proposes a new embedding for that gene tree. When a new species network is proposed, the rebuild-embedding operator proposes a new embedding for each gene tree in the species network. If there is no valid embedding for any gene tree, the gene tree or species network proposal is rejected.

The rebuild-embedding operator proposes a new embedding proportional to the product of traversal probabilities across all traversed reticulation nodes. Specifically, we defin_Q_e the (unnormalized) likelihood of a compatible embedding *x* as 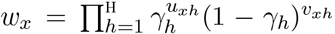, where h is the number of reticulation nodes in the species network, *u_xh_* is the number of lineages traversing node *H_h_* to the branch associated with *γ_h_*, and vxh is the number of lineages traversing node *H_h_* to the alternative branch associated with 1 − *γ_h_*. If there is no reticulation in the species network (i.e., it is a species tree), *w_x_* = 1. For example in Figure 2, there are two possible embeddings for one gene tree (orange) while the likelihoods are *γ*^2^ (current) and (1 − γ)^2^ respectively, and four possible embeddings for the other gene tree (blue) while the likelihoods are γ ^2^, γ (1 − γ), (1 − γ)γ, and (1 − γ)^2^ (current), respectively.

The proposal ratio of moving from embedding *x* to *x*' is

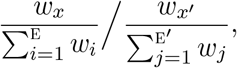

where E and E' are the number of possible embeddings in the current and new states respectively. If E' = 0 (no valid embedding), the move is aborted. This proposal distribution is chosen to have a superior acceptance ratio than if a new embedding is proposed randomly from all possible embeddings.

### 2.5 Summarizing posterior distribution of species networks

Reducing many hundreds of posterior or bootstrap samples to a summary result is essential in order to describe the underlying distribution. For phylogenetic trees, many summary methods have been developed such as “majority rule consensus” and “maximum clade credibility” trees (Heled and Bouckaert, 2013). By comparison, methods to summarize samples of phylogenetic networks are underdeveloped. As part of the `SpeciesNetwork` package, we have implemented a basic method for summarizing networks, where unique network topologies are reported in descending order of their posterior probabilities. For each unique topology, each subnetwork is annotated with its posterior probability and node age credible interval.

To facilitate the calculation of posterior probabilities and credible intervals, we have developed an algorithm to enumerate each unique subnetwork, and label all occurrences of a unique subnetwork in a sample of phylogenetic networks. After running this algorithm, the label of a network's root node uniquely identifies its topology, and the generation of a sorted summary of posterior topologies becomes trivial. Details of the algorithm are given in the Appendix. The default setting of our summary tool eliminates all parallel branches (e.g., *S*_3_ *H*_2_ in Fig. 5b) from all samples in the posterior before summarizing, which simplifies the posterior distribution of networks and reduces the number of unique topologies.

Alternatively, users may generate a summary network using the “major displayed tree” method as implemented in the PhyloNetworks package (Solís-Lemus et al., 2017).

## 3 Simulations

The components from the last section, i.e., the unnormalized posterior density and the operators, allow us to implement a Markov chain Monte Carlo (MCMC) procedure to sample species networks and gene trees from the posterior distribution, given a multilocus sequence alignment. The implementation is available within BEAST 2 (Bouckaert et al., 2014) as an add-on `SpeciesNetwork`. A convenient format for the species networks, and a link to our source code, is presented in the Appendix.

We investigate the performance of the implementation using simulations in this section. Time is measured by genetic distance (substitutions per site) throughout the simulations, so that *θ = Nμ* is used for all population sizes and *τ_i_ = t_i_ μ* for the time of node *i*. The substitution rate μ is fixed to 1.0 across all gene lineages (strict molecular clock) and all loci (no rate variation).

### 3.1 Simulation and MCMC sampling without sequence data

To verify the implementation of our Bayesian MCMC method, we compared stochastic simulation to MCMC sampling of species networks and gene trees. We first generated networks under the birth-hybridization process. The simulator starts from the time of origin (*t*_0_) with one species. A species splits into two (speciation) with rate λ, and two species merge into one (hybridization) with rate *v*. At the moment of *k* branches, the total rate of change is 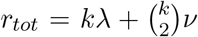. We generate a waiting time ~ exp(*r_tot_*) and a random variable *u ~ U*(0, 1). If *u < kλ/r_tot_*, we randomly select a branch to split; otherwise, we randomly select two branches to join, and generate an inheritance probability *γ ~ U*(0, 1). The simulator stops at time 0 (cf. Fig. 1). In this simulation, *τ*_0_ = 0.1, *λ* = 20, *v* = 10, and we kept 200,000 networks with exactly three tips. All the population sizes were fixed to *θ* = 0.01. Given each simulated species network, we then simulated a gene tree with two samples from each species (2, 2, 2) under the backward-in-time MSNC, resulting in 200,000 gene trees.

In the MCMC, we used all the operators for the species network (with 3 tips), gene tree (with 2 samples per species), and embedding (see above). The parameters *τ*_0_, *λ*, *v* and *θ* were fixed to the truth. The likelihood of data was set to be constant (no data). The chain was run 500 million steps and sampled every 2000 steps. The last 200,000 sampled species networks and gene trees were kept (i.e., the burn-in was 20%).

Theoretically, we expect the distributions of species network and gene trees to be identical from both simulation and MCMC sampling. Indeed, the networks obtained from the simulator and MCMC match when comparing the network length, root height, number of hybridizations, and time of the youngest hybridization (Fig. 6). The tree sets from MSNC and MCMC also give rise to the same distributions of tree length, the gamma-statistic (Pybus and Harvey, 2000), and Colless' index (Blum et al., 2006) as expected (Fig. 7).

**Figure 6:**
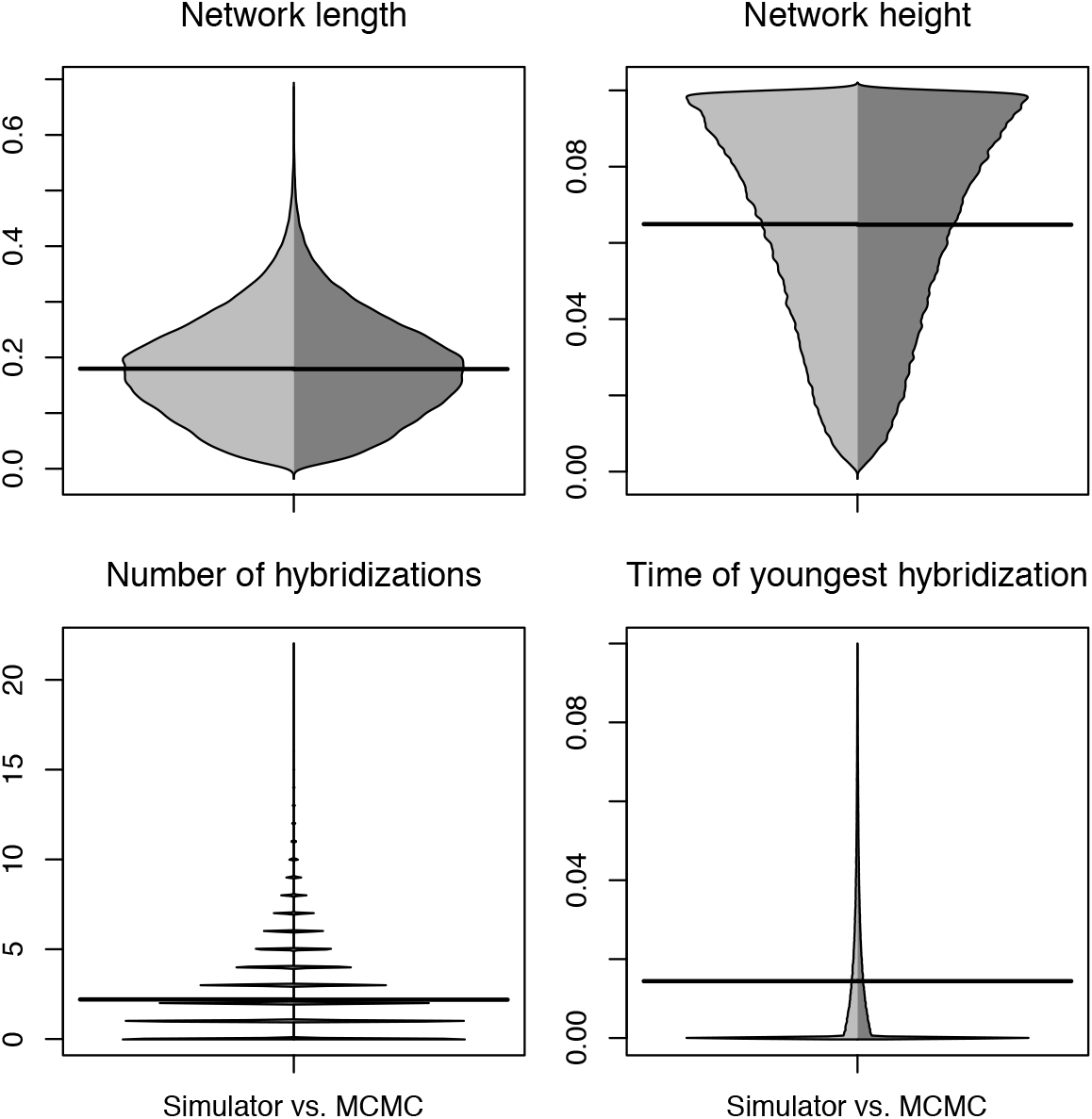
Beanplot of network summary statistics comparing 3-tips networks simulated under the birth-hybridization process (left, light gray) with those sampled using the MCMC operators (right, dark gray). The horizontal bar is the mean.

**Figure 7:**
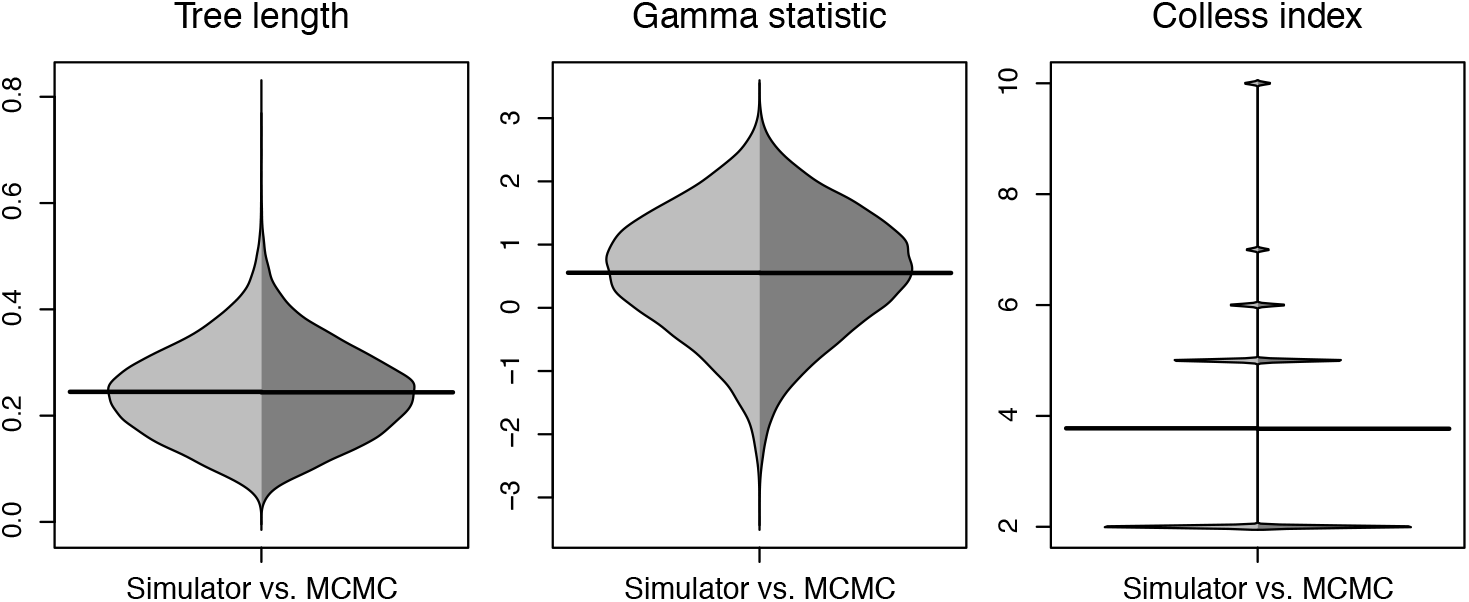
Beanplot of three tree summary statistics comparing gene trees simulated under MSNC (left, light gray) with those sampled using the MCMC operators (right, dark gray). The sample configuration was (2, 2, 2).

### 3.2 Inference of species networks from sequences

We simulated sequence alignments of multiple loci to reveal the ability of our method to recover the true species network from multilocus sequence data. The true network is shown in Figure 1a, with *τ*_1_ = 0.05, *τ*_2_ = 0.03, *τ*_3_ = 0.02, *τ*_4_ = 0.01, *γ* =0.3, and population sizes *θ* = 0.01. A random gene tree was generated for each locus under the MSNC. Then DNA sequences of length 200 bp were simulated under JC69 model (Jukes and Cantor, 1969) along each tree. The sample configurations were (2, 4, 2) (meaning species *A* has 2, *B* has 4, and *C* has 2 sampled sequences) and (5, 10, 5), and the number of loci was 2, 5, 10, 20, 40, respectively. Under each setting, the simulation was repeated 100 times. In the inference, the priors were *τ*_0_ ~ exp(10) with mean 0.1, *d = λ − v* ~ exp(0.1) with mean 10, *r = v/λ ~ U*(0, 1), and *γ ~ U*(0, 1). The population sizes were integrated out analytically using inverse-gamma(5, 0.05) (Eq. 4). The substitution model was set to JC69 (the true model). We fixed *μ* = 1.0 for all genes as in the simulation (strict molecular clock and no rate variation). The MCMC chain was run 50 million steps and sampled every 2000 steps. The first 35% samples were discarded as burn-in. The results are shown in Figure 8.

**Figure 8:**
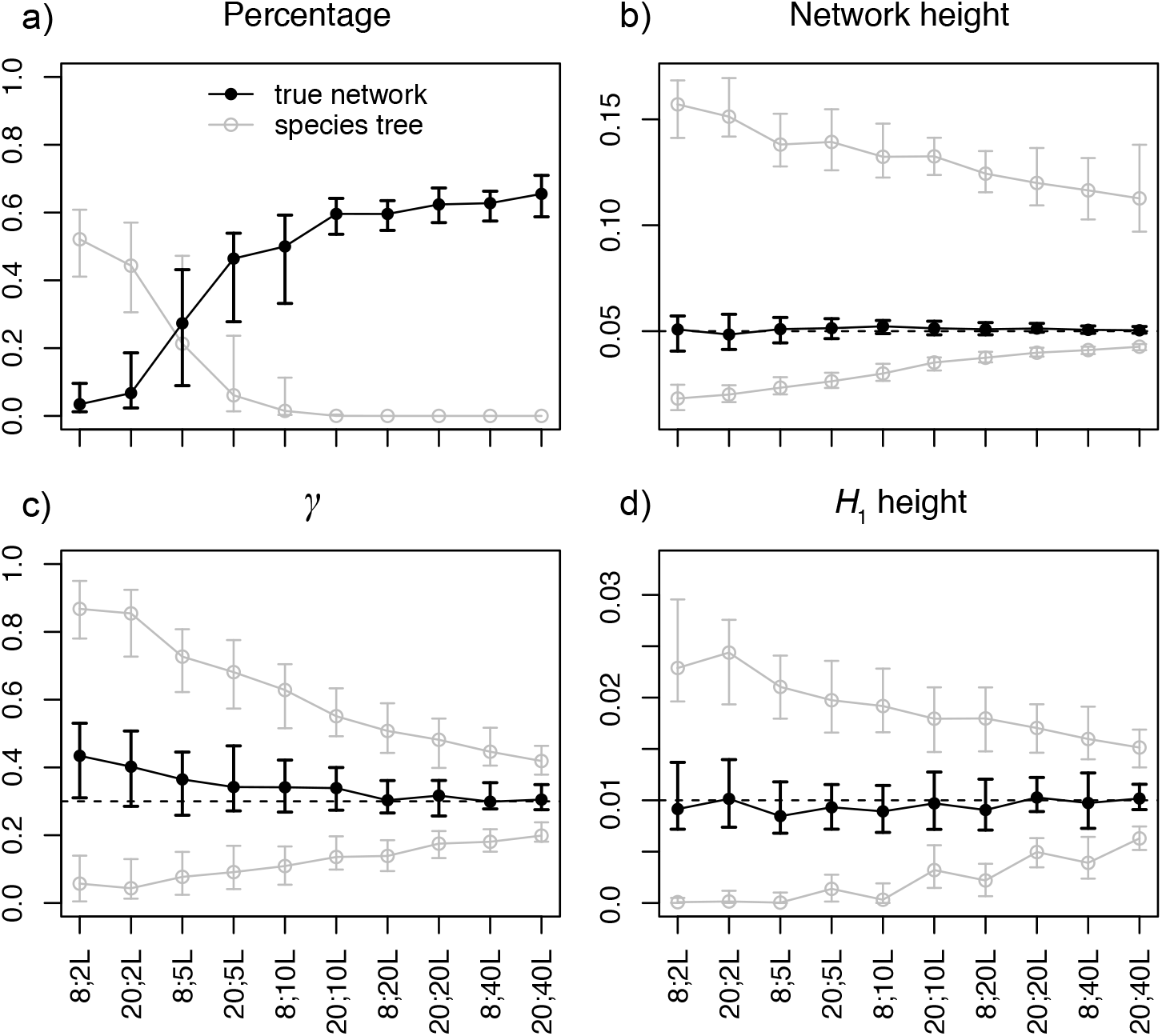
Posterior estimates of a) probability of the true network (black) and species trees (gray), b) network height, c) *γ* in the true network topology, and d) *H*_1_ height in the true network topology, when the data were simulated under the network in Figure 1a with sample configuration (2, 4, 2) or (5, 10, 5), and 2, 5, 10, 20, or 40 loci, respectively. For each setting in a), the dot/circle with error bars are the median and the 1^st^ and 3^rd^ quartiles of the percentages of 100 replicates. For each setting in b), c) and d), the black dot with error bars are the median and the 1^st^ and 3^rd^ quartiles of the posterior medians of 100 replicates, the gray circles with error bars are the same summaries for the 95% HPD intervals. The dashed lines indicate the true values.

With only 2 loci, the species trees are inferred with the highest posterior probability, the distribution of species network topologies is sensitive to the prior (Fig. 8a). The HPD intervals of the network height are also very wide (Fig. 8b). As sample size increases, the posterior estimates become increasingly accurate. Conditional on the true species network topology inferred (i.e., Fig. 1a), the estimates of inheritance probability *γ* and time of hybridization become increasingly accurate as the number of loci increases (Fig. 8cd). We also observe that adding more loci increases the accuracy of inference more than adding more individuals. For example, by comparing (5, 10, 5) 5 loci with (2, 4, 2) 10 loci, the latter has higher probability of recovering the true species network (Fig. 8a).

To reveal the power of our method to detect both ancient and recent hybridization events, we simulated gene trees and sequences subsequently under the true species network shown in Figure 1b, with *τ_R_* = 0.05, *τ*_*H*_1__ = 0.03, *γ*_1_ = 0.6, *τ*_*H*_2__ = 0.01, *γ*_2_ = 0.7, *τ*_*S*_1__ = 0.035, *τ*_*S*_2__ = 0.04, *τ*_*S*_3__ = 0.012, *τ*_*S*_4__ = 0.015, and population sizes *θ* = 0.01. The sample configurations were (2, 2, 2, 2) and (5, 5, 5, 5) respectively. The other settings were kept the same as in the previous simulation. The results are shown in Figure 9.

**Figure 9:**
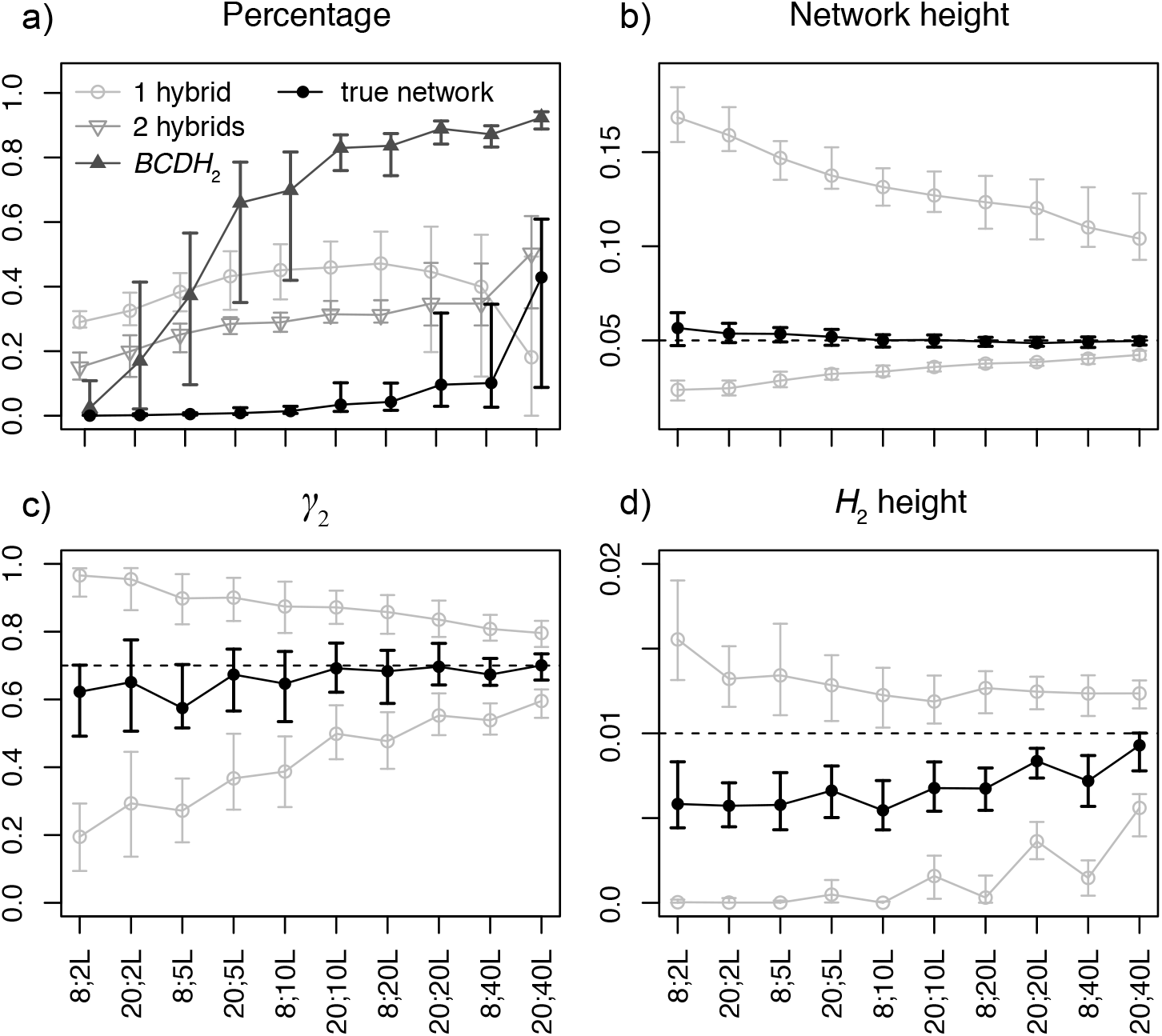
Posterior estimates of a) probability of the true network (black), networks with 1 or 2 hybridizations (light gray), and networks with the *BCDH*_2_ subnetwork (dark gray), b) network height, c) *γ*_2_ in the *BCDH*_2_ subnetwork, and d) *H*_2_ height in the BCDH2 subnetwork, when the data were simulated under the network in Figure 1b with sample configuration (2, 2, 2, 2) or (5, 5, 5, 5), and 2, 5, 10, 20, or 40 loci, respectively. For each setting in a), the dot/circle with error bars are the median and the 1^st^ and 3^rd^ quartiles of the percentages of 100 replicates. For each setting in b), c) and d), the black dot with error bars are the median and the 1^st^ and 3^rd^ quartiles of the posterior medians of 100 replicates, the gray circles with error bars are the same summaries for the 95% HPD intervals. The dashed lines indicate the true values.

We find that an ancient hybridization close to the root is much harder to detect than a recent hybridization. With up to 8 samples and 20 loci, the posterior probabilities of the true network topology are all close to zero. The estimates start to increase with 20 samples and 20 loci or more (Fig. 9a). The difficulty is mainly due to the fact that there are few gene-tree lineages close to the root of the network, making it hard to distinguish the true hybridization event with incomplete lineage sorting in the ancestral populations. However, the recent hybridization event is inferred with high probability using 10 to 40 loci (Fig. 9a). More specifically, we looked at the posterior probability of networks having the *BCDH*_2_ subnetwork structure (cf. Fig. 1b). Conditional on having this subnetwork inferred, the estimates of inheritance probability γ_2_ become increasingly accurate as the number of loci increases (Fig. 9c), although the time of hybridization *H*_2_ is generally underestimated (Fig. 9d). It is not feasible to perform larger scale simulations, e.g., using 100 loci or more, to investigate the power of recovering the ancient hybridization (thus the true species network). Further studies need to be carried out after the efficiency of the operators is improved (see Discussion).

## 4 Analysis of biological data

### 4.1 Three closely related spruce species

We analyzed a dataset of three spruce species (*Picea purpurea, P. likiangensis* and *P. wilsonii*) in the Qinghai-Tibet Plateau (Sun et al., 2014). *P. purpurea* was inferred to be a homoploid hybrid of *P. likiangensis* and *P. wilsonii* (Sun et al., 2014). The original data has 166 diploid individuals and 11 nuclear loci (50 from *P. wilsonii*, 56 from *P. purpurea*, 60 from *P. likiangensis*, and two phased haplotype sequences per individual per locus).

To achieve proper mixing and convergence in a reasonable time, the data was truncated into two non-overlapping datasets by randomly selecting individuals, resulting in 20 individuals from *P. purpurea*, 15 from *P. likiangensis*, and 15 from *P. wilsonii* (100 sequences per locus) for each. The priors for the species network were *τ*_0_ ~ exp(500) with mean 0.002, *d = λ − v* ~ exp(0.01) with mean 100, *r = v/λ ~ U*(0, 1), and *γ ~ U*(0, 1). The population sizes were integrated out analytically (Eq. 4) using inverse-gamma(3, 0.003) with mean 0.0015 and mode 0.00075. The substitution model was HKY85 (Hasegawa et al., 1985), with independent κ (transition-transversion rate ratio) and state frequencies at each locus. The clock rate was fixed to 1.0 (strict molecular clock across branches) and gene-rate multipliers were used to account for rate variation across loci. The MCMC chain was run for 1 billion steps and sampled every 20,000 steps. The first 35% of samples were discarded as burn-in. For each dataset we obtained two independent runs, and the two runs were checked for effective sample sizes (ESS) and the consistency of trace plots of inferred parameters. The MCMC samples from the two runs were combined.

The species network shown in Figure 10 has the highest posterior probabilities for the two datasets, both are > 0.95. This confirms that *P. purpurea* is a hybrid species of *P. likiangensis* and *P. wilsonii*. The estimates of γ are 0.33 (0.18, 0.52) and 0.37 (0.17, 0.57) respectively (median and 95% HPD interval). To investigate the impact of prior on population sizes, we fixed the species network topology to the one in Figure 10, and used three priors for the population size parameter: inverse-gamma(3, 0.0003) with mean 0.00015 (small), inverse-gamma(3, 0.003) with mean 0.0015 (medium), and inverse-gamma(3, 0.03) with mean and 0.015 (large), respectively. The population sizes were either inferred using MCMC or integrated out analytically. The other priors and MCMC settings were unchanged.

**Figure 10:**
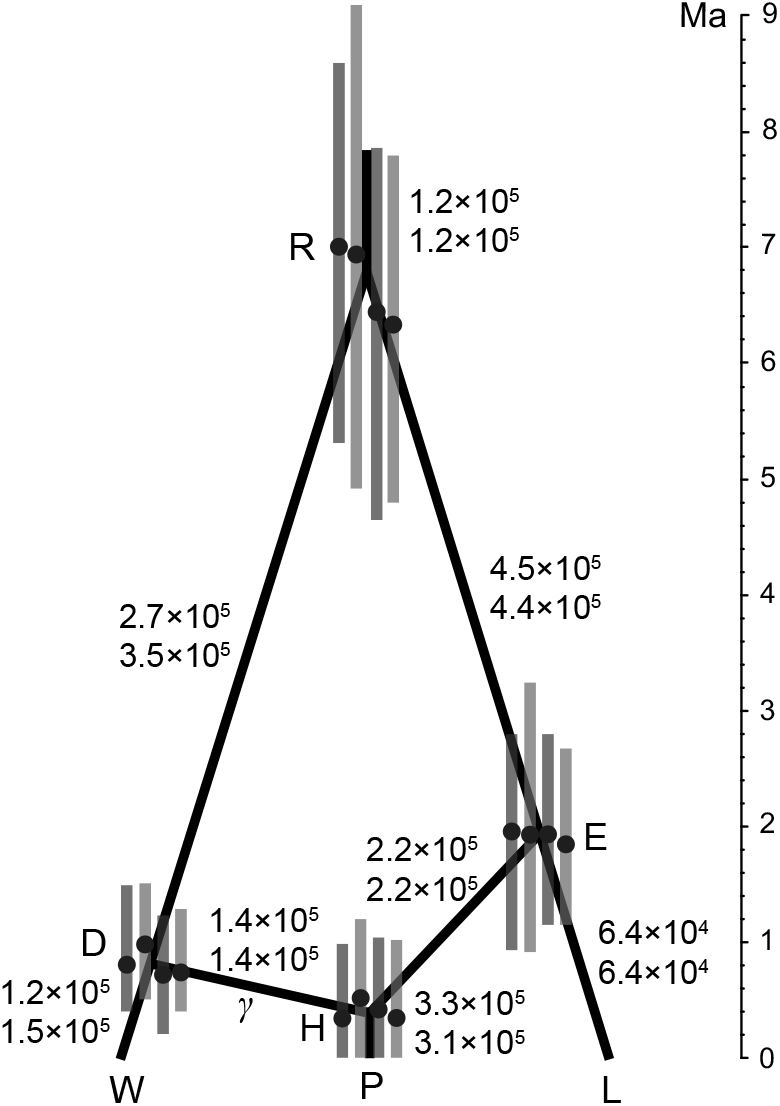
The species network with highest posterior probability (> 95%) inferred from the spruce data. The medians and 95% HPD intervals of node heights (black dots with error bars) are in unit of million years. From left to right, they are for dataset 1 with population sizes inferred and integrated out, and dataset 2 with population sizes inferred and integrated out, under the inverse-gamma(3, 0.003) prior. The numbers beside the branches are the medians of effective population sizes inferred from dataset 1 (above) and 2 (below). See also Table S1 and S2.

The posterior estimates of γ, node heights, and population sizes are summarized in Supplemental Table S1 and S2. The estimates are similar for both datasets regardless of whether the population sizes were inferred or integrated out under the same prior, but some estimates vary slightly across different priors. Below we summarize the results from the inverse-gamma(3, 0.003) prior (medium mean) for population sizes (Fig. 10, and middle column of Table S1 & S2). Around 31–37% of the nuclear genome of *P. purpurea* was derived from *P. wilsonii* (thus 63–69% from *P. likian*-gensis). This estimate is close to the original estimate of 31% made using approximate Bayesian computation (ABC) (Sun et al., 2014). Assuming an average substitution rate *μ* = 2 × 10^-4^ per site per million years (Sun et al., 2014), and dividing the node heights (τ's in Table S1 & S2) by *μ*, we get the times measured by million years (Fig. 10). The time of hybridization is inferred to be around 1 Ma. The estimate was 1.3 (0.73, 2.2) Ma in the original analysis assuming the same height for nodes *D, E*, and *H*. Moreover, we get an older and narrower estimate for the root age (Fig. 10), compared to 2.7 (1.4, 6.5) Ma in the original analysis. Similarly, dividing estimates of *θ*'s (Table S1 & S2) by *μ* =1 × 10^-8^ per site per generation, we get the effective population sizes (Fig. 10). The inferred population sizes of *P. purpurea*, *P. wilsonii*, and *P. likiangensis* are smaller than those estimated using ABC (cf. Table 4 in Sun et al., 2014).

### 4.2 Seven yeast species (Saccharomyces)

We re-analyzed another dataset of seven yeast species, including *S. cerevisiae* (Scer), *S. paradoxus* (Spar), *S. mikatae* (Smik), *S. kudriavzevii* (Skud), *S. bayanus* (Sbay), *S. castellii* (Scas), and *S. kluyveri* (Sklu). There are in total 106 orthologous gene loci and one sequence per species per locus (Rokas et al., 2003). This data analyzed using concatenation under maximum likelihood yielded a single tree (Fig. 11a) with 100% bootstrap values at every branch (Rokas et al., 2003). The analysis using BEST (Liu, 2008) showed two main species trees in the posterior (Fig. 11ab)(Edwards et al., 2007). Both studies discovered extensive incongruent phylogenies from individual genes, with phylogenetic conflict often involving Scas and Sklu. Recently, the full dataset was also analyzed using a Bayesian method co-estimating species networks and gene trees. Extensive hybridization events were found, usually involving Scas and Sklu as the donor or recipient (Wen and Nakhleh, 2017).

**Figure 11:**
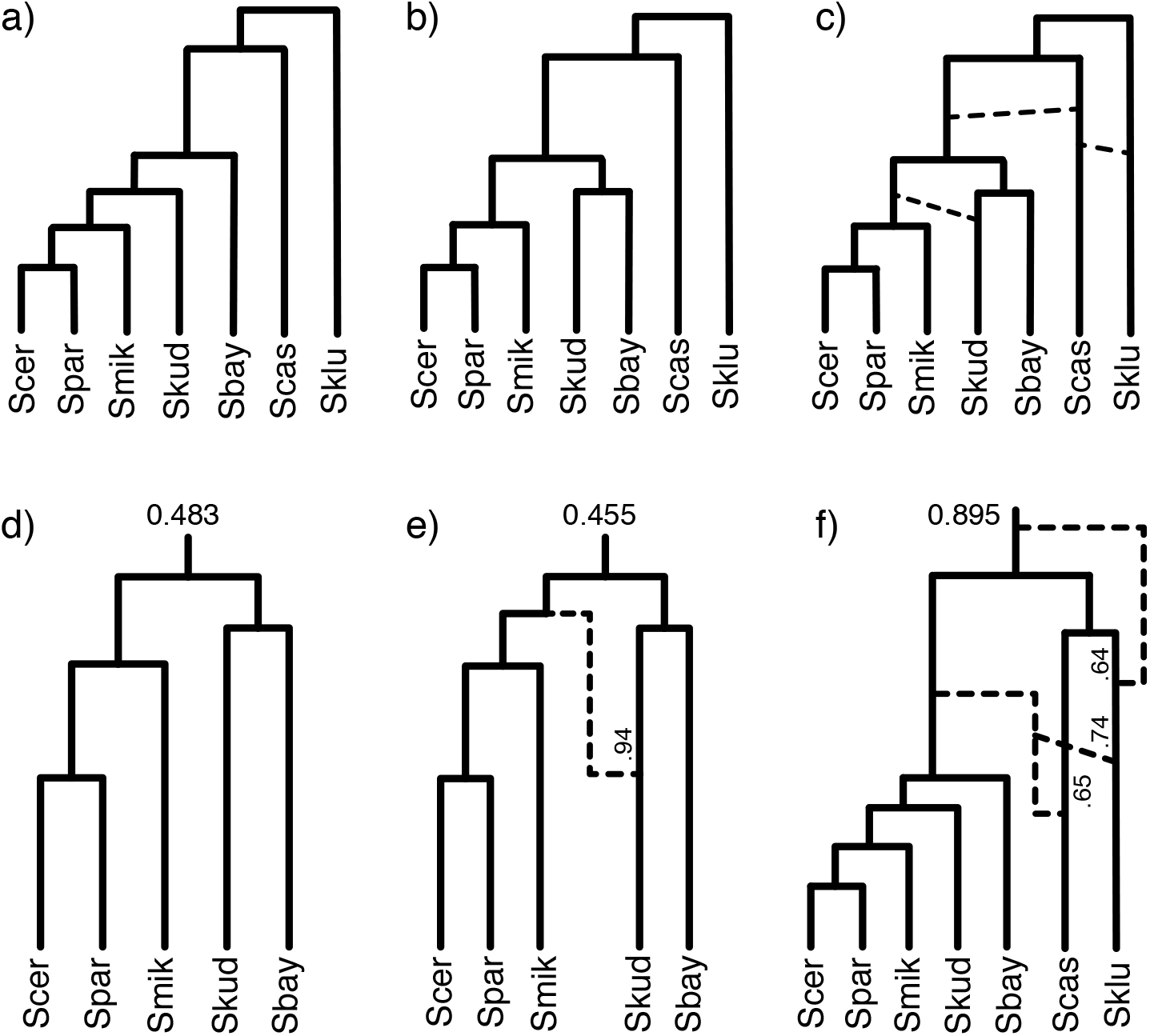
The species networks inferred from the yeast data. a) The species tree estimated using concatenation under maximum likelihood (Rokas et al., 2003). a) and b) are the two main species trees in the posterior analyzed using BEST (Edwards et al., 2007). c) The representative species network inferred using our method on all 7 species and 106 loci. The dashed lines indicate possible hybridization events. d) and e) are two species networks in the 95% posterior credible set using 5 species and 106 loci (excluding Scas and Sklu). f) The species network inferred using 7 species and 28 loci with strong phylogenetic signal. The posterior probabilities are labelled at the root. The larger inheritance probabilities are labelled beside the corresponding branches.

For the inference using our method, the priors for the species network were *τ*_0_ ~ exp(10) with mean 0.1, *d = λ − v* ~ exp(0.2) with mean 5, *r = v/λ ~ U*(0, 1), and *γ ~ U*(0, 1). The population sizes were integrated out analytically using inverse-gamma(3, 2θ), while the mean population sizes θ was assigned a gamma(2, 100) prior with mean 0.02. We still used the HKY85 substitution model (Hasegawa et al., 1985), gene-rate multipliers for rate variation across loci, and the same MCMC chain settings as in the spruce data analysis.

Similarly, we observed extensive hybridizations among Scas, Sklu, and the rest five species (Fig. 11c) in the posterior estimates from independent runs. The incongruence among gene tree phylogenies are well captured and explained by the hybridization events. These patterns are similar to the results in Wen and Nakhleh (2017). The backbone tree (by removing the reticulation branches with smaller inheritance probabilities from the networks) is consistent with the species tree in Figure 11b. However, the complexity of hybridizations caused difficulty and poor mixing of MCMC using the full data. The species network topology may stay unchanged for fairly long steps during the MCMC and independent runs give different number and direction of hybridizations, although the hybridization pattern and the backbone tree are the same across runs.

Using only five species by excluding Scas and Sklu produced consistent results across runs, and the posterior samples from the three runs are combined. About half of the samples in the 95% posterior credible set are specie trees (Fig. 11d) and another half are species networks with one hybridization leading to Skud (Fig. 11e). The result of Wen and Nakhleh (2017, Fig. 22e) showed only one species network in the 95% posterior credible set with opposite hybridization direction (from Skud to Sbay) and smaller γ then ours (0.75 vs. 0.94). But both analyses have the same backbone tree as in Figure 11d. The difference is probably due to the different priors and evolutionary models we used (see Discussion). The root heights are both 0.094 (0.092, 0.096) (median and 95% HPD interval) in Figure 11de. The branch lengths are measured by genetic distance (mean substitutions per site). The posterior estimate of mean population sizes *θ* is 0.00086 (0.00015, 0.0018). The rate multipliers range from 0.55 to 1.5 for the 106 loci.

We further investigated the 28 loci with strong phylogenetic signal, as done in Wen and Nakhleh (2017). The gene trees inferred from these loci under maximum likelihood have at least four internal branches with bootstrap support > 70%. The priors and MCMC settings are the same as for the 106 loci. Using all the seven species, the species network with highest posterior probability (0.895) is shown in Figure 11f. Three hybridization events are recovered, all involving Scas and Sklu. The recent two hybridizations were also found in Wen and Nakhleh (2017, Supplemental material) and the inheritance probabilities were similar to ours. In addition, we found an extra hybridization event after the early divergence of Scas and Sklu, older than the other two. When using only five species by excluding Scas and Sklu in a separate run, the species tree with the same topology as the subtree in Figure 11f was inferred with highest posterior probability (0.98). This is different from the backbone tree using all 106 loci (cf. Fig. 11d), indicating conflicting phylogenetic signal among loci.

## 5 Discussion

Methods to build a species network (e.g., Wu, 2010; Park et al., 2010; Albrecht et al., 2012) traditionally use inferred gene trees from each locus without accounting for their uncertainties, and employ nonparametric criteria such as parsimony. For population level data, the sequences are similar and the signal for gene tree topologies is typically low, so using fixed gene trees is assigning too much certainty to the data. These methods typically assume that gene tree discordance is solely due to reticulation, thus may suffer in the presence of incomplete lineage sorting (Yu et al., 2011). The MSNC model (Yu et al., 2014) provides a statistical framework to account for both incomplete lineage sorting and reticulate evolution. But properly analyzing genetic data to infer species networks under the MSNC model is a challenging task. There have been methods using only the gene tree topologies from multiple loci under MSNC (Yu et al., 2012, 2014; Wen et al., 2016). However, gene trees with branch lengths are more informative for inferring species tree or network topology than gene tree topologies alone. Accounting for branch lengths can improve distinguishability of species networks (Pardi and Scornavacca, 2015; Zhu and Degnan, 2017). Although methods using estimated gene trees (with branch lengths) from bootstrapping or posterior sample as input take account gene tree uncertainty (Yu et al., 2014; Wen et al., 2016), directly using sequence data to co-estimate species networks and gene trees in a Bayesian framework showed improved accuracy (Wen and Nakhleh, 2017). Pseudo-likelihood approaches (Yu and Nakhleh, 2015; Solís-Lemus and Ané, 2016) compute faster than full likelihood or Bayesian approaches, but have severe distinguishability issues and require more data to achieve good accuracy.

At the time of writing, another Bayesian method inferring species networks and gene trees simultaneously from multilocus sequence data was released (Wen and Nakhleh, 2017). The general framework here is similar, but we highlight four major differences. We use a birth-hybridization prior for the species network which naturally models the process of speciation and hybridization. The prior is extendable to account for extinction, incomplete sampling, and rate variation over time, as we outline below. Wen and Nakhleh (2017) used a descriptive prior combining a Poisson distributed number of reticulations with exponential distributed branch lengths. Secondly, we allow parallel branches in the network. This is biologically possible. Even if the true species history has no parallel branches, the observed species network can still have such features due to incomplete sampling. Note though that a very large number of individuals and loci are required to detect such parallel branches. To prevent the species network from growing arbitrarily big, such that it becomes indistinguishable by the gene trees (Pardi and Scornavacca, 2015; Zhu and Degnan, 2017), we typically assign an informative prior to ensure the hybridization rate is lower than the birth rate. A similar strategy was used in Wen et al. (2016); Wen and Nakhleh (2017) by restricting the rate of the Poisson distribution. Third, we account for the uncertainty in the embedding of a gene tree within a species network by estimating the MSNC probability conditional on a proposed embedding at each MCMC step. This provides a posterior distribution of gene trees and their embeddings within a species network, enabling analysis of which alleles are derived from which ancestral species. The cost instead is additional MCMC operations compared to integrating over all embeddings at each step (Wen et al., 2016; Wen and Nakhleh, 2017). Last but not least, we applied analytically integration for population sizes in the species network (Eq. 4). This reduces the number of parameters for the rjMCMC operators to deal with, and should improve convergence and mixing. Finally, our implementation in SpeciesNetwork is an extension to BEAST 2 (Bouckaert et al., 2014), to take advantage of many standard phylogenetic models, such as different substitution models, relaxed molecular clock models, and the BEAUTi graphical interface.

In our approach, we employ a simple prior for the species network based on a birth-hybridization model. Analogous to priors for species trees (e.g., Stadler, 2010; Heath et al., 2014), the prior for species network could be extended to account for speciation, extinction, hybridization, and incomplete sampling, each with a different rate, leading networks with present-day samples and potential past samples corresponding to fossils. The rates could also be allowed to vary over time, to model the diversification patterns during speciation (the skyline model for trees, Stadler et al., 2013). When considering networks instead of trees, techniques to derive the probability density of trees cannot be directly applied as the hybridization rate depends on pairs of lineages rather than individual lineages. This non-linearity necessitates solving differential equations to derive the species network probability densities, a task which we defer to a later study.

Our approach is limited in computational speed. The empirical analysis was done, e.g., on only three species with 50 individuals and 11 loci, or up to seven species and 106 loci but one individual per species. The main bottleneck is the MCMC operators. Due to hard constraints between the species network and embedded gene trees (Fig. 2), MCMC operators changing them separately limit the ability to analyze genomic scale data from many individuals. More specifically, updating the species network will likely violate a gene tree embedding, resulting in very low acceptance rate of the operator. Thus it will be essential to design more efficient MCMC operators. There have been coordinated operators that can change species tree and gene trees simultaneously (Rannala and Yang, 2003, 2017; Jones, 2017). Such operators are possible to be extended to species networks, and will potentially improve efficiency of the MCMC algorithm. Proposing new embeddings of gene trees in species network is also costing. Thus it might be worthwhile to integrate over the embeddings (Wen et al., 2016; Wen and Nakhleh, 2017) if they are not of interest. Moreover, there are methods to integrate out the gene trees under the multispecies coalescent model when analyzing biallelic genetic markers (RoyChoudhury et al., 2008; Bryant et al., 2012; Zhu et al., 2017). However, it is not yet feasible to apply this strategy to multilocus sequence alignment. Computationally, implementing Metropolis-coupled MCMC (MC^3^, Geyer, 1991) will help to overcome multiple local peaks in the posterior, and further parallelizing the cold and hot chains will gain speed.

In summary, we developed a Bayesian method for inferring species networks together with gene trees and evolutionary parameters from multilocus sequence data. The method is implemented within a general Bayesian framework, with potential future extensions to the theoretical model and to the practical implementation.

## Acknowledgments

This research was supported by the European Research Council under the Seventh Framework Programme of the European Commission (PhyPD: grant number 335529 to T.S.). C.Z. acknowledges his salary as well as a visit covered by this grant to the Centre for Computational Evolution, University of Auckland, New Zealand in mid-2016. We sincerely thank Simone Linz for detailed discussion on modeling phylogenetic networks.

## 6 Appendix

### 6.1 Numbering and labeling subnetworks across a sample

We describe an algorithm by pseudocode to enumerate all unique subnetwork topologies within a sample of phylogenetic networks. Apart from subnetwork topologies consisting of a single node (i.e. leaf nodes), each topology label has a corresponding set of child subnetwork topology numbers. The algorithm works by recursively constructing the mapping of parent to child subnetwork topology numbers, beginning at the root or origin node of each phylogenetic network.

~~~
Initialize the counter *i* to 0
Initialize the (node label set to node label) map m
~~~

~~~
For each taxon *t*:
Assign *i* as the label of *t*
Increment *i*
~~~

~~~
For each phylogenetic network *s*:
Begin Recursion from the oldest node of *s*
~~~

~~~
Recursion:
Input: A network node *n*
Output: A label *l* to identify the subnetwork topology of *n*
~~~

~~~
If *n* is a leaf node:
Get the label *l* of the taxon *t* of *n*
Else:
Initialize the node label set *d*
For each child node nc of *n*:
Get *l_c_* by continuing Recursion from *n_c_*
Add *l_c_* to *d*
~~~

~~~
If *d* is in *m*:
Get the label *l* of *d*
Else:
Set *l* to the value of *i*
Link *d* to *l* in *m*
Increment *i*
Return *l*
~~~

### 6.2 Proposing embeddings proportional to their likelihoods

We describe an algorithm by pseudocode to propose compatible gene tree embeddings, given a species network and a set of gene trees, in proportion to their embedding likelihoods. The algorithm works by stochastically constructing an embedding during a depth-first search of a gene tree. When a gene tree lineage traverses a bifurcation node, there is a set of compatible embedding histories (for the subtree defined by the gene tree branch) where the lineage descends through the left child branch of the bifurcation node, and another set for the right child branch. A left or right embedding is chosen at random weighted by the sum total of embedding likelihoods for each child branch of the bifurcation node, to ensure that embeddings are proposed in proportion to their likelihoods.

The likelihood for the proposed embedding is also computed during the depth-first search; when a gene tree lineage traverses a reticulation node, its likelihood is multiplied by the *γ_h_* or the (alternative) 1 − *γ_h_* probability. When a coalescent event occurs, the likelihoods of the left and right subtrees are multiplied. Because embeddings are proposed in proportion to their likelihoods, the MCMC proposal probability is the embedding likelihood normalized by the sum total of compatible embedding likelihoods.

~~~
Given the species network *s*:
For each gene tree *g*:
Get the root gene tree node *gtn_r_* from *g*
Get the root species network branch *snb_r_* from *s*
Try to get *e*, *l*, and *t* by Recursion from *gtn_r_* and *snb_r_*
~~~

~~~
If there is no compatible embedding:
Reject the proposal
Else:
Propose *e* as the new embedding
Multiply the proposal probability by (*l ÷ t*)
~~~

~~~
Recursion:
Input 1: A gene tree node *gtn*
Input 2: A species network branch *snb*
Output 1: An embedding *e*
Output 2: Its likelihood *l*
Output 3: The total likelihood *t*
~~~

~~~
If *gtn* traverses through the tipward node of *snb*:
For each child branch *snb_c_* of *snb*:
If there is any compatible embedding of *gtn* through *snb_c_*:
Get *e_c_*, *l_c_*, and *t_c_* by Recursion from *gtn* and *snb_c_*
Add the traversal of *gtn* through *snb_c_* to *e_c_*
If the tipward node of *snb* is a reticulation:
Multiply *l_c_* by *γ_h_* or 1 − *γ_h_*
Multiply *t_c_* by *γ_h_* or 1 − *γ_h_*
~~~

~~~
Pick one *snb_c_* at random weighted by *t_c_*
Set the embedding *e* to the value of *e_c_* for the chosen *snb_c_*
Set the likelihood *l* to the value of *l_c_* for the chosen *snb_c_*
Calculate the total likelihood *t* as the sum of all *t_c_*
Else:
If *gtn* is a leaf:
Initialize an embedding *e*
Initialize the likelihood *l* to 1
Initialize the total likelihood *t* to 1
Else:
For each child node *gtn_c_* of *gtn*:
Get *e_c_*, *l_c_*, and *t_c_* by Recursion from *gtn_c_* and *snb*
~~~

~~~
Construct the embedding *e* by merging both *e_c_*
Calculate the likelihood *l* as the product of both *l_c_*
Calculate the total likelihood *t* as the product of both *t_c_*
~~~

~~~
Return *e*, *l*, and *t*
~~~

### 6.3 Representation of phylogenetic networks

The species network can be represented using extended Newick format (Cardona et al., 2008), which was also used in the software PhyloNet (Than et al., 2008).

For example, the species network in Figure 1a is written as

~~~
((A:0.02,(B:0.01)#H1[&gamma=0.3]:0.01)S1:0.03,
(#H1:0.02,C:0.03)S2:0.02)R:0.03;
~~~

where the hash sign indicates a reticulation node, and the inheritance probability is in the brackets as metadata. Such extended Newick string can be read into IcyTree (icytree.org; Vaughan, 2017) and be displayed nicely.

### 6.4 Software availability

The method is implemented in the add-on SpeciesNetwork for BEAST 2 (Bouckaert et al., 2014), including the inference, simulation, and summary tools, and is hosted publicly on GitHub (https://github.com/zhangchicool/speciesnetwork).

